# Csp1, A Cold-Shock Protein Homolog in *Xylella fastidiosa* Influences Pili Formation, Stress Response, and Gene Expression

**DOI:** 10.1101/2021.07.21.453299

**Authors:** Wei Wei, Teresa Sawyer, Lindsey P. Burbank

**Affiliations:** USDA Agricultural Research Service, San Joaquin Valley Agricultural Sciences Center, Parlier, CA; Electron Microscopy Facility, Oregon State University, Corvallis, OR

## Abstract

Bacterial cold shock-domain proteins (CSPs) are conserved nucleic acid binding chaperones that play important roles in stress adaptation and pathogenesis. Csp1 is a temperature-independent cold shock protein homolog in Xylella fastidiosa, a bacterial plant pathogen of grapevine and other economically important crops. Csp1 contributes to stress tolerance and virulence in X. fastidiosa. However, besides general single stranded nucleic acid binding activity, little is known about the specific function(s) of this protein. To further investigate the role(s) of Csp1, we compared phenotypic differences between wild type and a csp1 deletion mutant (Δcsp1). We observed decreases in cellular aggregation and surface attachment with the Δcsp1 strain compared to the wild type. Transmission electron microscopy imaging revealed that Δcsp1 had reduced pili compared to the wild type and complemented strains. The Δcsp1 strain also showed reduced survival after long term growth, in vitro. Since Csp1 binds DNA and RNA, its influence on gene expression was also investigated. Long-read Nanopore RNA-Seq analysis of wild type and Δcsp1 revealed changes in expression of several genes important for attachment and biofilm formation in Δcsp1. One gene of intertest, pilA1, encodes a type IV pili subunit protein and was up regulated in Δcsp1. Deleting pilA1 increased surface attachment in vitro and reduced virulence in grapevines. X. fastidiosa virulence depends on bacterial attachment to host tissue and movement within and between xylem vessels. Our results show Csp1 may play a role in both virulence and stress tolerance by influencing expression of genes important for biofilm formation.

**Importance:** *Xylella fastidiosa* is a major threat to the worldwide agriculture industry (1, 2). Despite its global importance, many aspects of *X. fastidiosa* biology and pathogenicity are poorly understood. There are currently few effective solutions to suppress *X. fastidiosa* disease development or eliminate bacteria from infected plants(3). Recently, disease epidemics due to *X. fastidiosa* have greatly expanded(2, 4, 5), exacerbating the need for better disease prevention and control strategies. Our studies show that Csp1 is involved in *X. fastidiosa* virulence and stress tolerance. Understanding how Csp1 influences pathogenesis and bacteria survival can aide in developing novel pathogen and disease control strategies. We also streamlined a bioinformatics protocol to process and analyze long read Nanopore bacterial RNA-Seq data, which has previously not been reported for *X. fastidiosa*.

## Introduction

*Xylella fastidiosa* is an economically important plant pathogen that causes disease in many agricultural crops including grapevines, citrus, almonds, alfalfa, and coffee. Infection of grapevines by *X. fastidiosa* subsp. *fastidiosa* is known as Pierce’s disease(6). Pierce’s disease is a serious problem for the grapevine industry in the United States, especially in California where the disease threatens the $30 billion wine industry(7). During the infection cycle, *X. fastidiosa* is spread via sap feeding insect vectors and colonizes the xylem tissue of plants(8). In the plant xylem, the bacteria encounter many stressors such as plant defense responses that can reduce pathogen viability. Abiotic stressors such as cold temperature can also affect long-term survival of the bacteria in grapevines and has been linked to pathogen elimination and vine recovery(9, 10).

Due to its reduced genome size in comparison to other similar bacteria, *X. fastidiosa* lacks some well-developed stress responses, which can reduce cell viability and survival(11). One notable difference is the *X. fastidiosa* genome encodes only two known cold shock protein (Csp) homologs, Csp1 and Csp2(12), while other bacteria like *E. coli* and *Salmonella enterica* have upwards of nine(13–15). Most research on bacterial cold shock proteins have focused on their role in helping bacteria adapt and survive at suboptimal temperatures, however several studies show some Csps also contribute to virulence and general stress response(14, 16–18). In *E. coli*, five of the nine Csps (CspA, CspB, CspE, CspG and CspI) are induced by changes in temperature(15), while CspC and CspE are constitutively expressed at normal growth temperatures (37°C) and are involved in regulating stress response gene expression(13). *E. coli* CspD is induced at early stationary phase and is important for survival under nutrient poor conditions(18). Previous studies on *X. fastidiosa* revealed that Csp1 and Csp2 are not induced by cell exposure to cold conditions (12)(Burbank, unpublished). Data on Csp2 function(s) is limited because attempts to make *X. fastidiosa csp2* deletion mutants were unsuccessful. Deleting *csp1* resulted in reduced survival after cold treatment *in vitro,* however its importance to *X. fastidiosa* cold survival *in planta* is not as well established(12). Csp1 is also important for osmotic stress tolerance(12). These results suggest that like *E. coli* CspE and CspC, *X. fastidiosa* Csp1 may be less important for cold survival and play a more prominent role in general stress tolerance.

In some animal and plant pathogens, Csps are also important for regulation of virulence factors. A triple deletion mutant (Δ*cspABD*) in bacterial foodborne pathogen *Listeria monocytogenes* reduced oxidative and cold stress survival and impaired host cell invasion and intracellular growth(16). The *L. monocytogenes* Δ*cspABD* mutant was also deficient in cellular aggregation and did not express surface flagella or exhibit swarming motility(19). Gene expression analysis showed reduced expression of virulence and motility genes in *L. monocytogenes csp* mutants, suggesting some Csps may regulate gene expression. Similarly, some cold shock proteins in plant pathogenic bacteria also act as virulence factors and regulate gene expression. The *Xanthomonas oryzae pv. oryzae* (*Xoo*) CspA protein regulates expression of two virulence genes, *PXO_RS11830* and *PXO_RS01060*(17). Deletion of *Xoo cspA* decreased cold tolerance, bacterial pathogenicity, biofilm formation and polysaccharide production(17). In *X. fastidiosa* strain Stag’s Leap, deleting *csp1* resulted in significantly reduced disease severity and bacterial titer in the absence of cold stress in susceptible Chardonnay grapevines(12). However, the mechanisms of how Csp1 contributes to stress tolerance and virulence are not well understood.

In this study, we investigated the molecular mechanisms through which *X. fastidiosa* Csp1 contributes to stress tolerance and virulence. Since the *X. fastidiosa csp1* mutant was less tolerant to certain stress conditions and had lower bacterial titer compared to the wild type *in planta*(12), we compared long-term survival of wild type *X. fastidiosa* Stag’s Leap with the Δ*csp1* mutant and a complemented strain. We also investigated whether Csp1 influences biofilm formation since xylem occlusion by biofilms is a major aspect of *X. fastidiosa* pathogenicity, and the Δ*csp1* mutant produced less severe symptoms in grapevine compared to the wild type(12). Lastly, because Csp1 has general nucleic acid binding activity(12) and studies in other bacteria show some Csps regulate gene expression, we also investigated the influence of Csp1 on *X. fastidiosa* gene expression using RNA-Seq to compare transcriptomes of wild type Stag’s Leap and the Δ*csp1* mutant.

## Results

### *X. fastidiosa* Δ*csp1* mutant showed reduced long-term survival *in vitro*

Previous work showed that deleting the *csp1* gene in *X. fastidiosa* strain Stag’s Leap resulted in reduced tolerance to salt and cold stress(12). Since these results strongly suggest Csp1 may play a role in *X. fastidiosa* stress adaptation, we were interested if Csp1 was involved in survival under other stresses, such as prolonged growth times. We compared cell viability of wild type, Δ*csp1*, and Δ*csp1/csp1*+ strains grown on PD3 plates at 7 days post inoculation (DPI) when *X. fastidiosa* cells begin to decline, and 13 DPI (extended growth period). No significant change in viability was observed between the mutant and wild-type at 7 days post inoculation (DPI) (Figure 1A), but there was a significant decrease in viability of Δ*csp1* at 13 DPI compared to the wild-type and complemented strains (Figure 1B). These results suggest Csp1 is important for long term survival of *X. fastidiosa in vitro*.

**Figure 1.**
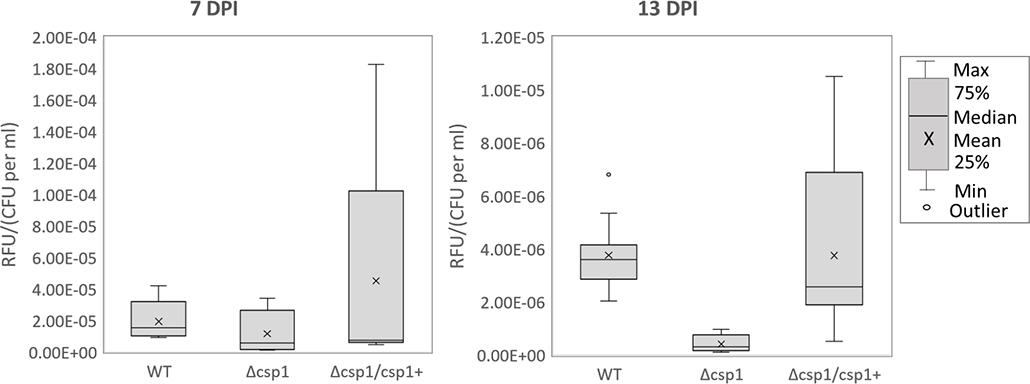
Cell Viability of the Δ*csp1* during long-term growth. Wild type Stag’s Leap, Δ*csp1*, and Δ*csp1*/*csp1*+ were grown on PD3 plates for up to 13 days. Cell viability was quantified at 7 days post inoculation (DPI) and 13 DPI using AlamarBlue (Life Technologies) fluorescent cell viability reagent by measuring RFU of each sample and normalizing to total cells quantified by qPCR. Graph represents data collected from at least three independent experiments. **Indicates treatment significantly different from the wild type based on one-way ANOVA followed by Tukey means comparison test (p<0.01).

### Δ*csp1* strain showed reduced cellular aggregation and surface adhesion

Biofilm formation is an important aspect of *X. fastidiosa* host colonization and involves both cell-cell aggregation and cellular adhesion to surfaces(20). Wild type Stag’s Leap cells form visible aggregates and a biofilm ring at the air-liquid interface when grown in liquid PD3 media (Figure 2A). However, the Δ*csp1* mutant showed a dispersed phenotype with visibly less cells attached at the air-liquid interface when grown under the same conditions as WT and the complemented strains (Figure 2A). We quantified and compared aggregation of the WT, Δ*csp1*, and Δ*csp1/csp1*+ strains and found that the percentage of aggregated cells in liquid culture was significantly lower for *Δcsp1* compared to the WT and complemented strains (Fig. 2B). In addition to cellular aggregation, biofilm formation also requires cell adhesion to surfaces. We quantified surface attachment of static cultures of WT, Δ*csp1*, and Δ*csp1/csp1*+ strains grown in 96-well plates using crystal violet staining and observed a significant decrease in the amount of attached cells for the mutant strain compared to WT and the complemented strains (Figure 2C).

**Figure 2.**
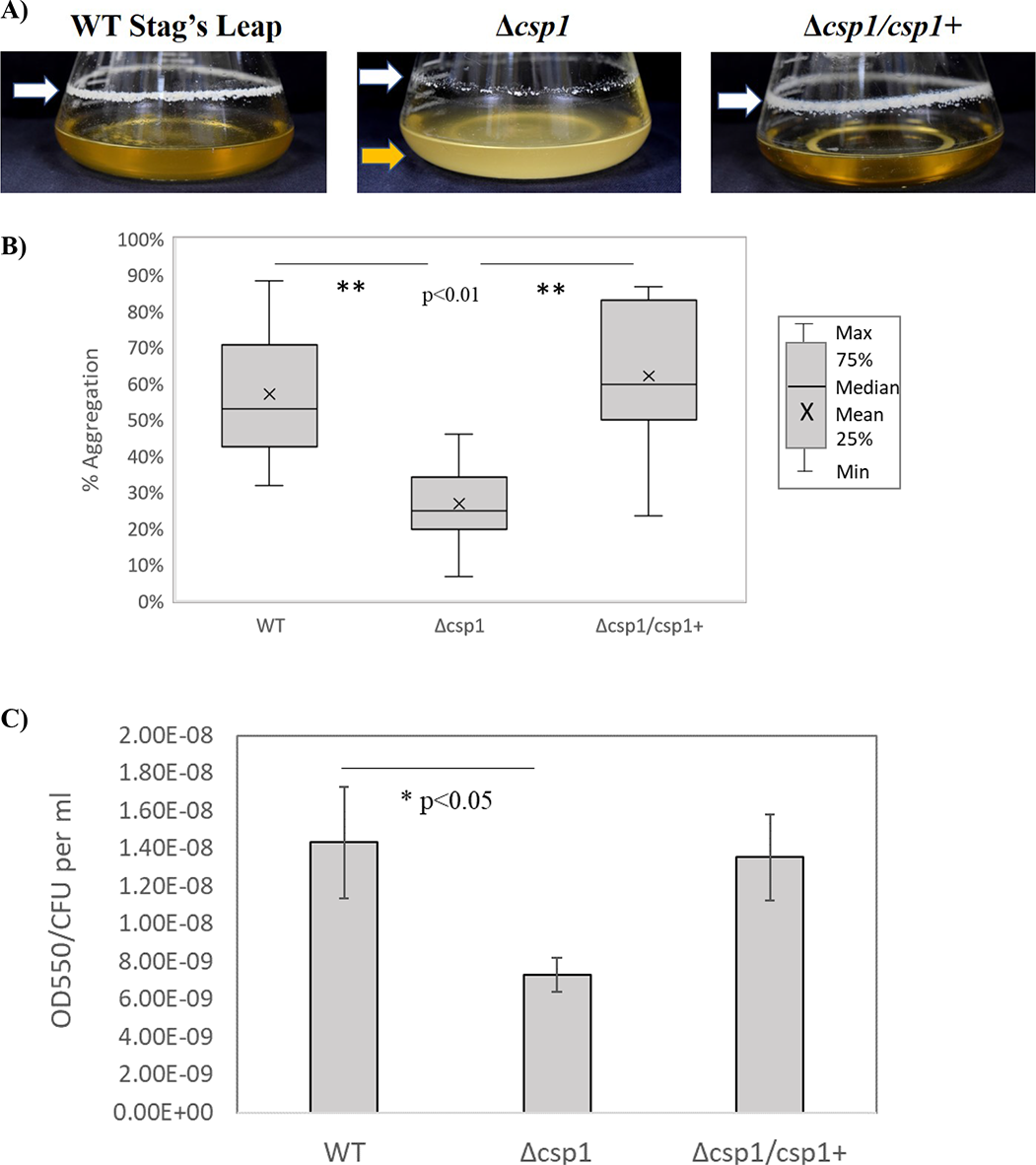
Cellular aggregation and attachment of Δ*csp1*, *in vitro.* **A)** Cell-cell aggregation and surface attachment was documented after 4 days of growth in liquid PD3 medium at 28°C with shaking at 180 rpm. The yellow arrow indicates the dispersed phenotype of the Δ*csp1* strain, and the white arrows indicate the ring of attached cells at the air-liquid interface. **B)** Cellular aggregation was quantified by measuring the OD600 of statically grown liquid cultures of WT, Δ*csp1*, and Δ*csp1/csp1*+ before and after manual dispersal of cells using the equation: [(OD600D-OD600U )/ OD600D]*100 where OD600D = optical density of dispersed culture and OD600U = optical density of undispersed culture. The graph represents a total of at least nine replicates from three separate experiments. **Indicates significant difference based on one-way ANOVA followed by Tukey means comparison test (p<0.01). **C)** Cell attachment was quantified by measuring the amount of crystal violet stain retained by cells attached to the walls of 96-well plates (OD550) after static growth for 4 days. OD550 was normalized to total cells (CFU/ml quantified by qPCR). Graph represents at least 45 technical replicates from three separate experiments. *Indicates significant difference (p<0.05) from wild type based on one-way ANOVA followed by Bonferroni-Holm.

### Δ*csp1* mutant shows reduced pili formation

*X. fastidiosa* pili are important for cellular aggregation and surface attachment(21). To evaluate the effect of *csp1* on pilus formation, cells of the wild type Stag’s Leap, Δ*csp1* mutant, and the complemented strains were visualized using Transmission electron microscopy (TEM), as shown in Figure 3. TEM images of wild-type *X. fastidiosa* shows pili localized to one pole of the cell, while the *csp1* mutant did not show visible pili formation (Figure 3). The complemented strain shows restored pilus formation, with most of the pili concentrated towards one pole of the cell (Figure 3). TEM was performed at Oregon State University’s Electron Microscopy Facility (Corvallis, OR).

**Figure 3.**
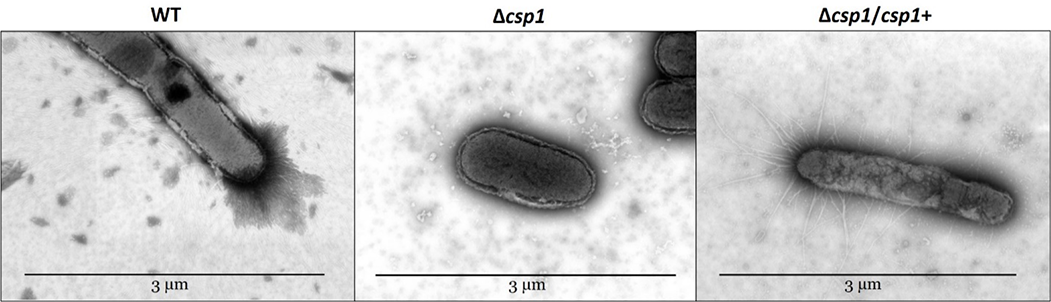
TEM images of *X. fastidiosa* strains. Pili location and abundance were observed for wild type Stag’s Leap (WT), the *csp1* mutant (Δ*csp1*), and the complemented (Δ*csp1/*Δ*csp1+*) strains using the Helios NanoLab 650 microscope.

### Motility and attachment related genes were differentially expressed in the Δ*csp1* strain

To investigate the influence of Csp1 on *X. fastidiosa* gene expression we used Nanopore RNA-Seq to sequence transcriptomes of wild-type *X. fastidiosa* strain Stag’s Leap and the Δ*csp1* deletion mutant under standard growth conditions (28°C). RNA-Seq analysis revealed 90 genes were differentially expressed in the Δ*csp1* strain compared to the wild type (Table 1). Of the 90 differentially expressed genes, 65 were down regulated and 25 were up regulated in Δ*csp1* compared to the wild type. The RNA-Seq results also showed that no transcripts mapped to the *csp1* gene in the mutant strain, verifying *csp1* was absent and showing the accuracy of transcript mapping (Table 1). Gene ontology analysis using GSEA Pro (http://gseapro.molgenrug.nl/ ) showed significant enrichment of genes involved in cell adhesion/attachment, gene regulation, translation, and rRNA binding. To identify genes of interest for further investigation, we focused on genes that may be associated with phenotypes observed in the Δ*csp1* mutant, such as reduced cellular aggregation and attachment. Several differentially expressed genes encoding proteins involved in attachment, motility and/or biofilm formation included *pilA1* (PD1924), *pilA2* (PD1926), *fimA* (PD0062), *fimC* (PD0061), *hsf*/*xadA* (PD0824), *pilV* (PD0020), and *fimT* (PD1735). We performed qRT-PCR to verify the Nanopore expression data for several of these genes and saw that expression of *pilA1* was consistently up regulated in the *csp1* mutant strain in all samples tested (Supplemental Table 1). Based on these results and findings from other studies showing *pilA1* is involved in *X. fastidiosa* attachment and biofilm formation(22), we chose to investigate the roles(s) of *X. fastidiosa* Stag’s Leap *pilA1* further.

**Table 1.** Differentially Expressed Genes

|  |  |  | Transcripts per million |  |  |
| --- | --- | --- | --- | --- | --- |
| Locus ID | Gene symbol | Product | WT | $\Delta csp1$ | Expression Ratio ($\Delta csp1$/WT) |
| PD0020 | *pilV* | pre-pilin leader sequence | 180.08 | 0.00 | 0.00 |
| PD0141 | *fabG* | 3-ketoacyl-(acyl-carrier-protein) reductase | 127.23 | 0.00 | 0.00 |
| PD1354 |  | hypothetical protein | 2582.82 | 0.00 | 0.00 |
| PD1380 | *csp1* | cold shock protein | 5290.81 | 0.00 | 0.00 |
| PD1701 | *dnaB* | replicative DNA helicase | 44.29 | 0.00 | 0.00 |
| PD1735 | *fimT* | type 4 fimbrial biogenesis protein | 131.92 | 0.00 | 0.00 |
| PD1944 | *rpsR* | 30S ribosomal protein S18 | 8011.60 | 0.00 | 0.00 |
| PD2095 |  | hypothetical protein | 119.23 | 0.00 | 0.00 |
| PD1926 | *pilA2* | fimbrial protein | 2184.57 | 66.44 | 0.03 |
| PD0216 | *cvaC* | colicin V precursor | 10631.85 | 842.10 | 0.08 |
| PD1931 | *sucD* | succinyl-CoA synthetase subunit alpha | 133.48 | 11.68 | 0.09 |
| PD1317 |  | hypothetical protein | 112.39 | 10.34 | 0.09 |
| PD1905 | *xrvA* | virulence regulator | 492.63 | 46.72 | 0.09 |
| PD2003 | *rplJ* | 50S ribosomal protein L10 | 479.16 | 45.72 | 0.10 |
| PD0463 |  | hypothetical protein | 2108.37 | 201.98 | 0.10 |
| PD0084 | *rplS* | 50S ribosomal protein L19 | 947.68 | 115.89 | 0.12 |
| PD1945 | *rpsF* | 30S ribosomal protein S6 | 1201.60 | 179.24 | 0.15 |
| PD1063 |  | hypothetical protein | 1287.41 | 202.47 | 0.16 |
| PD0556 |  | hypothetical protein | 28667.31 | 4555.29 | 0.16 |
| PD0062 | *fimA* | fimbrial subunit precursor | 10009.57 | 1724.44 | 0.17 |
| PD0217 |  | hypothetical protein | 2815.96 | 511.29 | 0.18 |
| Locus ID | Gene symbol | Product | WT | Δcsp1 | Expression Ratio (Δcsp1/WT) |
| PD1440 | *rpsT* | 30S ribosomal protein S20 | 6060.28 | 1246.99 | 0.21 |
| PD0708 |  | virulence regulator | 672.31 | 141.02 | 0.21 |
| PD1914 | *rpmI* | 50S ribosomal protein L35 | 21603.77 | 4619.53 | 0.21 |
| PD0626 | *ssb* | single-stranded DNA-binding protein | 484.98 | 103.71 | 0.21 |
| PD0061 | *fimC* | chaperone protein precursor | 477.40 | 108.95 | 0.23 |
| PD1087 |  | hypothetical protein | 2361.29 | 585.51 | 0.25 |
| PD0283 | *dksA* | DnaK suppressor | 927.35 | 238.22 | 0.26 |
| PD0313 | *pspB* | serine protease | 76.39 | 19.88 | 0.26 |
| PD0459 | *rpsK* | 30S ribosomal protein S11 | 2578.30 | 684.46 | 0.27 |
| PD0460 | *rpsD* | 30S ribosomal protein S4 | 949.37 | 255.44 | 0.27 |
| PD1913 | *rplT* | 50S ribosomal protein L20 | 2372.81 | 665.13 | 0.28 |
| PD0447 | *rplN* | 50S ribosomal protein L14 | 2169.26 | 615.27 | 0.28 |
| PD2122 | *rnpA* | ribonuclease P | 1479.05 | 424.04 | 0.29 |
| PD2121 | *yidC* | putative inner membrane protein translocase component YidC | 96.10 | 30.18 | 0.31 |
| PD0453 | *rplR* | 50S ribosomal protein L18 | 1681.26 | 541.74 | 0.32 |
| PD1684 |  | hypothetical protein | 15592.41 | 5202.73 | 0.33 |
| PD0462 | *rplQ* | 50S ribosomal protein L17 | 1407.81 | 481.26 | 0.34 |
| PD0458 | *rpsM* | 30S ribosomal protein S13 | 2889.36 | 991.26 | 0.34 |
| PD1557 | *apbE* | thiamine biosynthesis lipoprotein ApbE precursor | 132.21 | 46.45 | 0.35 |
| PD1807 | *ompW* | outer membrane protein | 3923.03 | 1396.21 | 0.36 |
| PD0442 | *rplV* | 50S ribosomal protein L22 | 2512.05 | 900.32 | 0.36 |
| PD0856 | *dcp* | peptidyl-dipeptidase | 111.54 | 42.17 | 0.38 |
| PD1808 |  | hypothetical protein | 35966.82 | 13605.11 | 0.38 |
| PD0824 | *hsf/xadA* | afimbrial adhesin surface protein | 43.10 | 16.51 | 0.38 |
| PD0464 | *comM* | competence-related protein | 257.79 | 98.85 | 0.38 |
| PD1993 | *csp2* | temperature acclimation protein B | 97359.93 | 38394.08 | 0.39 |
| PD0246 | *secG* | preprotein translocase subunit SecG | 1557.96 | 650.10 | 0.42 |
| PD0436 | *rpsJ* | 30S ribosomal protein S10 | 7969.22 | 3538.88 | 0.44 |
| PD1984 | *gacA* | transcriptional regulator | 694.76 | 310.36 | 0.45 |
| PD0448 | *rplX* | 50S ribosomal protein L24 | 2529.90 | 1134.00 | 0.45 |
| PD1558 | *comE* | DNA transport competence protein | 16765.37 | 7555.53 | 0.45 |
| PD1709 | *mopB* | outer membrane protein | 597.52 | 309.74 | 0.52 |
| PD2123 | *rpmH* | 50S ribosomal protein L34 | 36412.20 | 19544.34 | 0.54 |
| PD0446 | *rpsQ* | 30S ribosomal protein S17 | 11515.54 | 6378.49 | 0.55 |
| PD0451 | *rpsH* | 30S ribosomal protein S8 | 3394.73 | 1983.09 | 0.58 |
| Locus ID | Gene symbol | Product | WT | Δcsp1 | Expression Ratio (Δcsp1/WT) |
| PD0060 | *fimD* | outer membrane usher protein precursor | 75.88 | 46.68 | 0.62 |
| PD0452 | *rplF* | 50S ribosomal protein L6 | 1627.21 | 1022.85 | 0.63 |
| PD0461 | *rpoA* | DNA-directed RNA polymerase subunit alpha | 477.69 | 301.63 | 0.63 |
| PD0159 |  | hypothetical protein | 1722.76 | 1188.73 | 0.69 |
| PD1506 |  | hemolysin-type calcium binding protein | 44.91 | 33.05 | 0.74 |
| PD0443 | *rpsC* | 30S ribosomal protein S3 | 1071.77 | 882.49 | 0.82 |
| PD0437 | *rplC* | 50S ribosomal protein L3 | 902.22 | 780.56 | 0.87 |
| PD2001 | *rpoB* | DNA-directed RNA polymerase subunit beta | 132.27 | 117.96 | 0.89 |
| PD0444 | *rplP* | 50S ribosomal protein L16 | 2534.48 | 2298.26 | 0.91 |
| PD1467 |  | hypothetical protein | 33.00 | 104.77 | 3.18 |
| PD0887 | *ruvA* | Holliday junction DNA helicase RuvA | 185.24 | 613.78 | 3.31 |
| PD0718 | *nodQ* | bifunctional sulfate adenylyltransferase subunit 1/adenylylsulfate kinase protein | 96.67 | 322.35 | 3.33 |
| PD1589 | *btuB* | TonB-dependent receptor | 21.48 | 86.50 | 4.03 |
| PD0179 |  | hypothetical protein | 21.64 | 92.00 | 4.25 |
| PD1652 | *recB* | exodeoxyribonuclease V beta chain | 4.63 | 19.88 | 4.29 |
| PD1167 | *ugd* | UDP-glucose dehydrogenase | 41.30 | 177.84 | 4.31 |
| PD1924 | *pilA1* | fimbrial protein | 202.45 | 892.96 | 4.41 |
| PD1702 |  | hypothetical protein | 80.27 | 362.95 | 4.52 |
| PD0744 | *hsf* | surface protein | 22.05 | 114.75 | 5.20 |
| PD1829 | *xylA* | family 3 glycoside hydrolase | 4.24 | 22.78 | 5.37 |
| PD1703 |  | hypothetical protein | 82.67 | 492.01 | 5.95 |
| PD0405 | *rpfG* | response regulator | 44.40 | 271.61 | 6.12 |
| PD0292 | *argE* | acetylornithine deacetylase | 13.64 | 89.63 | 6.57 |
| PD1517 |  | hypothetical protein | 52.15 | 349.41 | 6.70 |
| PD1280 | *hspA* | low molecular weight heat shock protein | 832.27 | 5951.42 | 7.15 |
| PD1409 | *grx* | glutaredoxin-like protein | 76.86 | 559.19 | 7.28 |
| PD0521 |  | hypothetical protein | 946.01 | 7042.88 | 7.44 |
| PD1850 |  | M20/M25/M40 family peptidase | 13.43 | 107.39 | 7.99 |
| PD1531 |  | hypothetical protein | 538.53 | 5090.18 | 9.45 |
| PD1468 | *bolA* | morphogene BolA protein | 236.76 | 2306.64 | 9.74 |
| Locus ID | Gene symbol | Product | WT | $\Delta csp1$ | Expression Ratio ($\Delta csp1$/WT) |
| PD1392 | *gumF* | GumF protein | 13.10 | 233.62 | 17.83 |
| PD1222 |  | hypothetical protein | 321.58 | 12183.51 | 37.89 |
| PD0657 |  | hypothetical protein | 70.31 | 2857.66 | 40.65 |
| PD0215 | *cvaC* | colicin V precursor | 1211.81 | 105871.34 | 87.37 |

### Stag’s Leap *ΔpilA1* mutant showed increased biofilm formation and decreased cellular aggregation *in vitro*

*X. fastidiosa* PilA1 is a type IV pili subunit protein that contributes to biofilm formation(22). Studies in *X. fastidiosa* strains TemeculaL and WM1-1 showed deleting *pilA1* leads to overabundance of type IV pili and increased attachment and biofilm formation(22). We observed increased expression of *pilA1* and decreased cell-to-cell adhesion and decreased attachment to surfaces in the Stag’s Leap Δ*csp1* mutant, so we created a *pilA1* deletion mutant in the Stag’s Leap background to investigate whether PilA1 is involved in attachment and biofilm formation in this strain as well. Similar to Temecula L, the Stag’s Leap Δ*pilA1* strain showed an increase in surface attachment compared to the wild type and complemented strains (Figure 4A). The Δ*pilA1* mutant also had reduced cellular aggregation and a dispersed phenotype in liquid media (Figure 4B). TEM images of Stag’s Leap Δ*pilA1* show that like the TemeculaL Δ*pilA1* mutant, deleting *pilA1* in Stag’s Leap also lead to overabundance of pili distributed around the entire cell (Supplemental Figure S2). These results show that *pilA1* is important for attachment and biofilm formation in Stag’s Leap and suggests that the decrease in surface attachment observed in the Δ*csp1* mutant may be a result of increased expression of *pilA1*.

**Figure 4.**
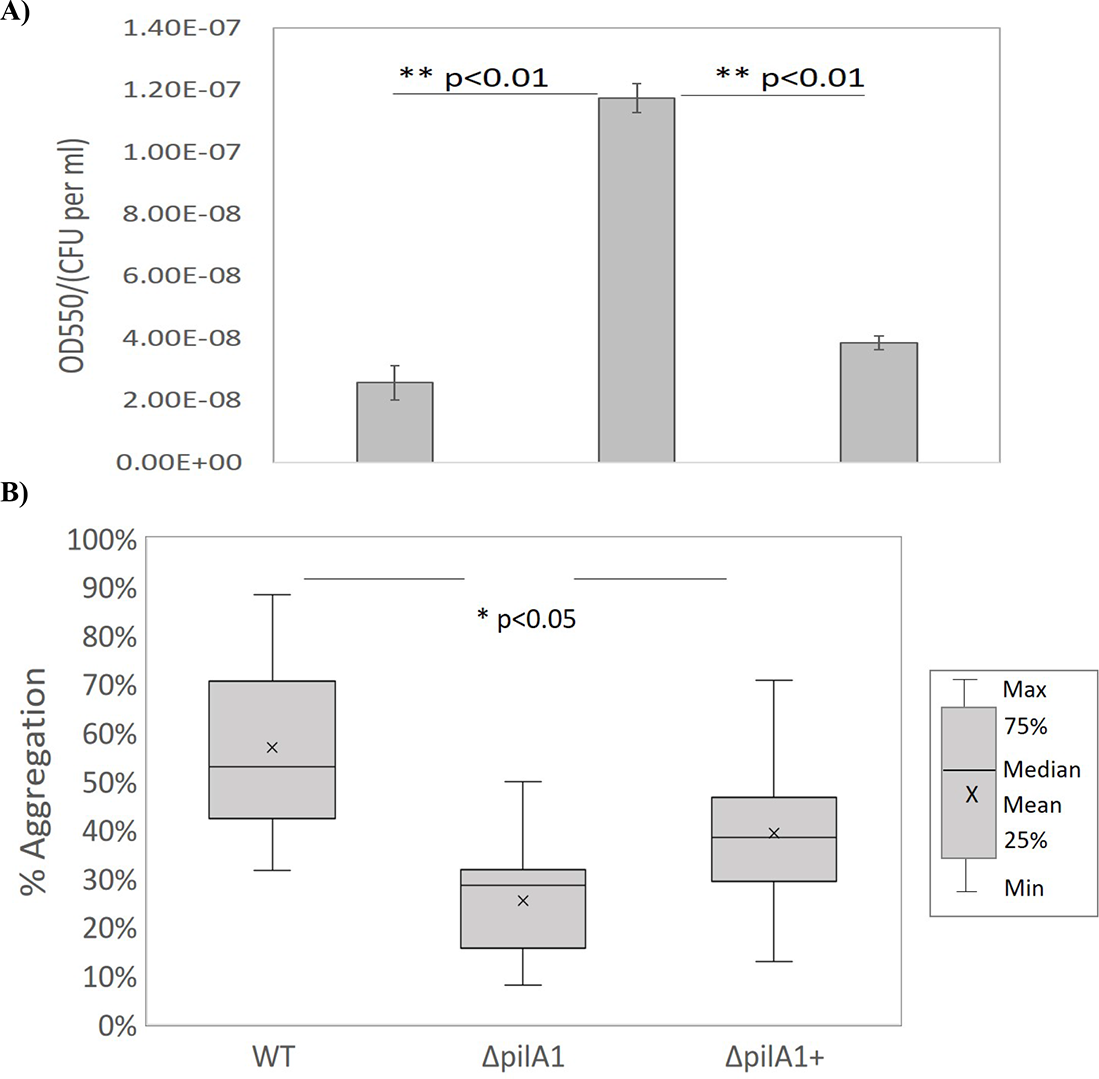
Cellular aggregation and attachment of Δ*pilA1.* **A)** Cell attachment of wild type Stag’s Leap (WT), *pilA1* deletion mutant (Δ*pilA1*)and *pilA1* complemented strain (Δ*pilA1*/*pilA1*+) was quantified by measuring the amount of crystal violet stain retained by cells attached to the walls 15ml polystyrene culture tubes (OD550) after static growth in 5ml of PD3 liquid media for 7 days. OD550 was normalized to total cells (CFU per ml) quantified by qPCR. Graph represents at least 16 technical replicates from four separate experiments. **Indicates significant difference from wild type based on one-way ANOVA followed by Tukey means comparison test (p<0.01). **B)** Cellular aggregation was quantified by measuring the OD600 of statically grown liquid cultures of WT, Δ*pilA1*, and Δ*pilA1/pilA1*+ before and after manual dispersal of cells. The percentage of aggregated cells was calculated using the equation from Figure 2B. The graph represents a total of at least 9 technical replicates from three separate experiments. *Indicates significant difference based on one-way ANOVA followed by Tukey means comparison test (p<0.05).

### The *ΔpilA1* mutant showed reduced virulence in grapevines

Xylem vessel occlusion caused by biofilms is one major mechanism of *X. fastidiosa* pathogenicity. Previous studies showed the Δ*csp1* mutant is less virulent in Chardonnay grapevines compared to wild type Stag’s Leap(12). Since both the Δ*csp1* and Δ*pilA1* mutant strains have altered biofilm phenotypes, and *pilA1* expression is up regulated in the Δ*csp1* strain, we were interested if *pilA1* also influences virulence in grapevines. We inoculated susceptible one-year-old potted Chardonnay grapevines with wild type Stag’s Leap, Δ*pilA1*, Δ*pilA1/pilA1*+ cultures or 1XPBS as the negative control. Disease severity was significantly reduced for plants inoculated with the *ΔpilA1* strain compared with plants inoculated with wild type and complemented strains at 16 weeks post-inoculation (Fig. 5A). The bacterial populations in inoculated plants were also quantified using qPCR. There was no significant difference in *X. fastidiosa* populations detected in petioles from plants inoculated with *ΔpilA1*, wild type, or the complemented strains at 16 weeks post-inoculation (Fig. 5B). This suggests that the virulence defect of the mutant strain was not due to reduced bacterial populations. Grapevine virulence assays were repeated the following year using the same *X. fastidiosa* strains and disease severity was calculated at 11 weeks post inoculation (11 weeks was used instead of 16 weeks due to time restraints). There was no significant difference in disease severity of grapevines inoculated with the different *X. fastidiosa* strains or the negative control at 11 weeks post inoculation (Supplemental Figure S3).

**Figure 5.**
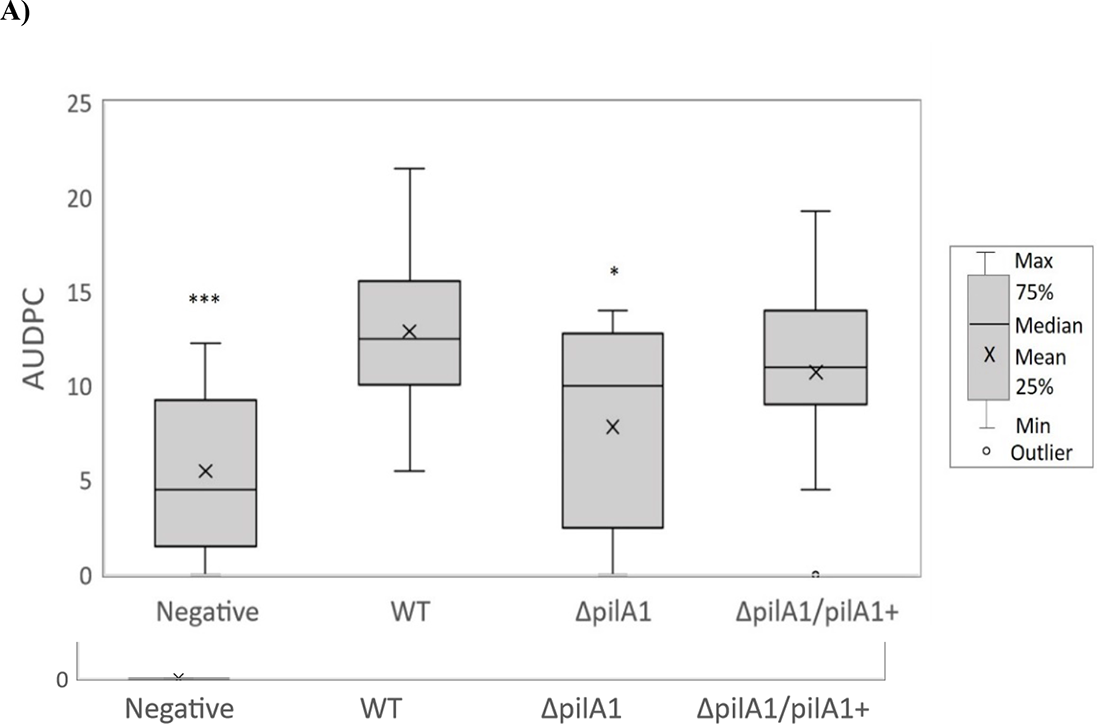
*ΔpilA1* has reduced symptom development in grapevines. One-year old potted grapevines (cv Chardonnay) were mechanically inoculated with wild type Stag’s Leap (WT), *pilA1* deletion mutant (Δ*pilA1*), the complemented (Δ*pilA1/pilA1*+) strain, or 1XPBS as the negative control. **A)** Symptom development in grapevines was scored on a 0-5 scale with 0 = no symptoms and 5 = plant death, over the period of 5-16 weeks post-inoculation. Area under the disease progress curve (AUDPC) was calculated using the Agricolae package for R. **B)** Bacterial populations in plant tissue were quantified using qPCR after 16 weeks post-inoculation and normalized to total DNA concentration. Graphs represents data from 20 plants inoculated with wild type, 15 plants inoculated with Δ*pilA1*, 15 plants inoculated with Δ*pilA1/pilA1*+, and 10 negative control plants. *Indicates treatment significantly different from the wild type based on one-way ANOVA followed by Tukey means comparison test (*= p<0.05, **= p<0.01, ***=p<0.001).

## Discussion

Bacterial Csp homologs are involved in a wide range of functions including cold tolerance, general stress response, and virulence(12, 17, 23). Csps involved in both stress response and virulence have been found in many animal pathogens such as *L. monocytogenes*(16, 19), *Brucella melitensis*(24)*, Salmonella enterica* serovar Typhimurium(14), *Enterococcus fecalis*(23), *Staphylococcus aureus*(25), as well as plant pathogens *Xanthomonas oryzae pv. oryzae*(17) and *X. fastidiosa*(12). Deleting Csp genes in these bacteria resulted in attenuated virulence, sometimes together with changes to cellular aggregation, surface attachment, and biofilm formation(19). However, the functional mechanisms underlying how Csps influence stress response and virulence are not well understood. In this study, we showed that *X. fastidiosa* Csp1 may have a role in regulating gene expression since deleting the *csp1* gene in strain Stag’s Leap influenced expression of 90 genes, several of which encode proteins important for bacteria attachment and motility such as the type IV pili gene *pilA1*. The phenotype of the Δ*csp1* mutant supported our transcriptome data, showing changes in cellular aggregation and surface attachment, processes that involve type I and type IV pili(21, 26). TEM images of Δ*csp1* showed a significant reduction in the number of visible pili compared to the wild type and complemented strains. Deleting *csp1* also reduced bacterial viability during late stationary phase, suggesting Csp1 is important for long-term survival.

Biofilm formation is an important part of *X. fastidiosa* insect vector and plant colonization(27). Xylem blockage by biofilms restricts water flow and is a major mechanism of *X. fastidiosa* pathogenicity(28). Biofilm development in *X. fastidiosa* is highly regulated and requires surface attachment and cellular aggregation, which are dependent on type I and type IV pili(29). Transcriptome analysis showed decreased expression of type I pilin gene *fimA* and type IV pili genes *pilA2* and *fimT*, while the type IV pili gene *pilA1* was up regulated in the Δ*csp1* mutant. The functions of *fimA*, *pilA2* and *fimT* were not investigated further in this study due to inconsistent gene expression results using quantitative RT-PCR to validate the RNA-Seq data. Other studies showed *X. fastidiosa* mutants lacking *fimA* had aggregation- and biofilm-deficient phenotypes, but were twitching-enhanced(26, 30), while mutants in *pilA2* were twitching-deficient compared to the wild type(22). Twitching motility allows *X. fastidiosa* to move against xylem flow and colonize tissue beyond the point of inoculation, making it important for virulence and plant colonization(31). Twitching motility of the Δ*csp1* mutant was not examined in this study but would be of interest for future studies to elucidate whether Csp1 influences this type of movement in *X. fastidiosa* and whether or not it contributed to the decreased virulence and bacterial titer observed in this strain. In addition, future experiments investigating *in planta* biofilm formation of the Δ*csp1* mutant can give us a better understanding of how Csp1 influences *X. fastidiosa* plant colonization.

We further investigated the role(s) of the type IV pilin protein PilA1 since qRT-PCR gene analysis for *pilA1* showed consistent up regulation of this gene in the Δ*csp1* strain in all samples tested. Type IV pilin proteins are flexible, filamentous structures mainly composed of a single pilin protein subunit encoded by *pilA* gene(s)(29). Genome analysis of *X. fastidiosa* showed the presence of at least four paralogs of the *pilA* gene(22). Most studies on bacterial type IV pili highlight their importance for twitching motility, however *pilA1* appears to be more involved in surface attachment and biofilm formation(22). When *pilA1* was deleted from the Stag’s Leap genome, we observed increased surface attachment and decreased cellular aggregation. Studies in *X. fastidiosa* strain TemeculaL and WM1-1 also showed deleting *pilA1* increased biofilm formation compared to the wild type strain(22). Electron microscopy of the Stag’s Leap, TemeculaL(22), and WM1-1(22) Δ*pilA1* mutants showed increased pili abundance with pili distributed around the entire cell, unlike their respective wild type strains which only showed pili localized to one cell pole. In contrast, the Δ*csp1* mutant, which showed up regulation of *pilA1*, appears to be deficient in pili formation compared to wild type Stag’s Leap.

The lack of visible pili in Δ*csp1* may contribute to the reduced attachment phenotype observed for this strain, while the increased abundance of type IV pili in the *ΔpilA1* mutants may contribute to increased attachment. Our Stag’s Leap *ΔpilA1* mutant was also less virulent in susceptible Chardonnay grapevines. *X. fastidiosa* disease symptom development is strongly correlated with pathogen spread within infected plants(21, 31), so increased surface attachment of the Δ*pilA1* mutant may restrict bacterial spread within the xylem, leading to the reduced virulence phenotype observed. However, there was no significant difference in bacterial titer between Δ*pilA1* and the wild type or complemented strains, suggesting the virulence defect of Δ*pilA1* may not be entirely due to reduced colonization. The *Δcsp1* mutant showed up-regulation of *pilA1* and decreased virulence in grapevines, while the *ΔpilA1* mutant also had reduced virulence in grapevines, indicating that other factors besides increased expression of *pilA1* are contributing to the Csp1-related virulence defect. Other variables that may be affecting virulence include decreased expression of virulence regulators (PD1905/*xrvA* and PD0708) in Δ*csp1*, or reduction in stress survival *in planta*. The functions of XrvA and the putative PD0708 protein in *X. fastidiosa* are still unclear but would be of interest to investigate in the future.

The Stag’s Leap Δ*csp1* mutant was less viable at 13 days post inoculation, which is considered late stationary phase of growth for this strain of *X. fastidiosa*, compared to the wild type and complemented strains. Bacteria in stationary phase encounter many stressors including nutrient limitation, accumulation of toxic by-products, and changes in pH, temperature, osmolarity, etc(32). Studies in other bacteria show that several temperature-independent cold shock proteins are involved in stationary phase stress response. *E. coli* CspD, which is 55.2% identical to the *X. fastidiosa* Csp1 amino acid sequence, is expressed during stationary phase upon glucose starvation and oxidative stress(18). The function of CspD is inhibition of DNA replication by nonspecific binding to single-stranded DNA regions at replication forks(33), and deletion of *cspD* leads to deceased persister cell formation while overexpression of *cspD* is lethal in *E. coli*(33). Bacterial persister cells are more resistant to antibiotics and can often be found in biofilm communities(34). Bacteria in biofilms are more resistance to host defense responses and antimicrobial compounds(35, 36), and have increased nutrient availability(37). Copper-based products are often used to control bacterial pathogens in agriculture, and transcriptome studies show that treating *X. fastidiosa* subsp*. pauca* biofilms with copper resulted in up-regulation of genes important for biofilm and persister cell formation including the toxin-antitoxin system MqsR/MqsA^29^ which in *E. coli*, regulates expression of *cspD.* The *E. coli* MqsR toxin is also directly involved in biofilm development and is linked to the development of persister cells(38). Overexpression of the *X. fastidiosa* MqsR toxin in a citrus pathogenic strain led to increased biofilm formation and decreased cell movement, resulting in reduced pathogenicity in citrus plants. In *X. fastidiosa* Temecula-1, an *mqsR* deletion mutant had reduced biofilm formation(39). MqsR over production also increased persister cell formation under copper stress in *X. fastidiosa*(40). It is unknown whether *csp1* expression in Stag’s Leap is regulated or influenced by the MqsR/MqsA complex, however functional similarities between Csp1 and CspD and the results from studies in *E. coli* showing *cspD* is directly regulated by MqsR/MqsA suggest this is a possibility. Future studies looking at possible links between Csp1 and the MqsR/MqsA toxin-antitoxin system can shed more light on *X. fastidiosa* stress tolerance and survival.

In summary, Csp1 is important for virulence and stress response in *X. fastidiosa*. Based on data from this study and past studies in *X. fastidiosa* and other bacteria, cold shock proteins like Csp1 may affect both pathogenicity and stress tolerance by influencing expression of genes important for biofilm formation. Biofilm formation is an essential virulence factor for *X. fastidiosa* and contributes to bacterial stress tolerance. The results of this study highlight the complexity of *X. fastidiosa* pathogen biology and more work looking at how cold shock proteins affect these processes will help us better understand how this pathogen colonizes and causes disease in hosts.

## Methods

### Bacterial culture conditions

he wild type strain used in this study is *Xylella fastidiosa* subspecies *fastidiosa* strain ‘Stag’s Leap’ isolated from grapevines with Pierce’s Disease in California, USA (41). The Δ*csp1* mutant strain used in this study has the *csp1* (PD1380) gene deleted and replaced with a Chloramphenicol resistance cassette(12). For all *in vitro* experiments, *X. fastidiosa* strains were grown on PD3(42) agar plates or liquid PD3 media without antibiotics or supplemented with 5 µg/mL of chloramphenicol and/or gentamycin when needed. *Escherichia coli* strains used for cloning and propagating of plasmid constructs were grown on LB medium supplemented with appropriate antibiotics at the following concentrations: chloramphenicol 35 µg/mL, spectinomycin 100 µg/mL, and gentamycin 10 µg/mL. All bacterial strains and plasmids used in this study are listed in Table 2 and 3, respectively.

**Table 2.** Bacterial strains.

| Strains | Description | Source |
| --- | --- | --- |
| *Xylella fastidiosa* subspecies *fastidiosa* Stag's Leap | Wild type strain, used to create mutant | (41) |
| *Xf* $\Delta csp1$ | *X. fastidiosa* deletion mutant in *csp1* (PD1380), Cm<sup>R</sup> | (12) |
| *Xf* $\Delta csp1/csp1+$ | Complemented strain, constructed by chromosomal insertion of *csp1* ORF at neutral site, Cm<sup>R</sup>, Gm<sup>R</sup> | (12) |
| *Xf* $\Delta pilA1$ | *X. fastidiosa* deletion mutant in *pilA1* (PD1924), Cm<sup>R</sup> | This study |
| *Xf* $\Delta pilA1/pilA1+$ | Complemented strain, constructed by chromosomal insertion of *pilA1* ORF at neutral site, Cm<sup>R</sup>, Gm<sup>R</sup> | This study |
| One Shot® TOP10 Chemically Competent *E. coli* | Commercially available *E. coli* strain used for propagating plasmid constructs, genotype: F- *mcrA* $\Delta(mrr-hsdRMS-mcrBC)$ $\Phi 80lacZ\Delta M15$ $\Delta lacX74$ *recA1* *araD139* $\Delta(araleu)7697$ *galU* *galK* *rpsL* (Str<sup>R</sup>) *endA1* *nupG* | Invitrogen |

### Construction of mutant and complemented strains

The *X. fastidiosa ΔpilA1* mutant strain was constructed by replacing the *pilA1* open reading frame with the Chloramphenicol resistance cassette from plasmid pCR8-csp1-chl(12) using homologous recombination. 931 bp of the upstream flanking region of the *pilA1* coding sequence was amplified from the WT Stag’s Leap gDNA using the primers pilA1-up-F/pilA1-up-R-SacI (Table 4). The pilA1-up-R-SacI adds a SacI restriction site to the 3’ end of the PCR product. 1.3kb of the downstream flanking region of the *pilA1* coding sequence was amplified using primers pilA1-down-F-XbaI/pilA1-down-R (Table 4). The pilA1-down-F-XbaI primer adds the XbaI restriction site to the 5’ end of the PCR product. The chloramphenicol resistance cassette from pCR8-csp1-chl(12) was amplified using primers Chl-F-SacI/Chl-R-XbaI, which adds SacI and XbaI restriction sites to the 5’ and 3’ ends, respectively, of the Chloramphenicol resistance cassette amplicon. All PCR reactions were performed using the high-fidelity Platinum™ *Taq* DNA Polymerase (Thermo Fisher). The chloramphenicol resistance cassette amplicon was ligated to the *pilA1* upstream and downstream flanking region amplicons using restriction enzyme cloning with SacI and XbaI (New England Biolabs) and T4 DNA ligase (Invitrogen). The ∼3.5kb ligation product was cloned into rapid TA cloning vector pCR8/GW/TOPO (Thermo Fisher) following the manufacturer’s instructions to create pCR8-ΔpilA1-chl. The TA cloning reaction was transformed into *E.coli* OneShot Top 10 competent cells (Thermo Fisher) and cells were spread onto LB agar supplemented with 100 µg/mL spectinomycin for selection of transformants. Transformants were screened for the correct 3.5kb insert using colony PCR with the primers pilA1-up-F/pilA1-down-R. Five colonies with the correct sized insert were inoculated into liquid LB supplemented with spectinomycin and chloramphenicol and grown overnight at 37°C for plasmid extraction using a QIAprep Spin Miniprep Kit (Qiagen). The plasmid constructs were confirmed by Sanger sequencing. Plasmid pCR8-ΔpilA1-chl was then transformed into WT *X. fastidiosa* using the natural transformation protocol(43) for mutagenesis via homologous recombination. Transformants were selected on PD3 agar supplemented with chloramphenicol and resistant colonies were screened by colony PCR using *X. fastidiosa* specific primers RST31/RST33(44) and gene specific primers to confirm the size of the insertion region (pilA1-ORF-F/pilA1-ORF-R, Table 4 ). The deletion mutation was confirmed by Sanger sequencing.

For complementation of the *pilA1* deletion, the *pilA1* ORF plus upstream and down stream flanking regions was inserted into the chromosome of the *X. fastidiosa* Δ*pilA1* strain at a neutral site as described(45). The *pilA1* ORF plus 405 bp of upstream and 215 bp of downstream sequence was PCR amplified from WT Stag’ Leap gDNA template using Platinum Taq polymerase and primers pilA1-OFR-405-F/pilA1-ORF-R (Table 4) and TA cloned into pCR8/GW/TOPO to created pCR8-pilA1-ORF (Table 3). Plasmid pCR8-pilA1-ORF was recombined with plasmid pAX1-GW(12) (Table 3) using the Gateway LR recombination protocol (Invitrogen). The resulting plasmid, pAX1-pilA-ORF, was purified from *E. coli* transformants and the correct insertion was confirmed by Sanger sequencing. pAX1-pilA-ORF was naturally transformed into the *X. fastidiosa ΔpilA1* strain, and transformants were selected on PD3 agar plates supplemented with gentamycin. Transformants were screened using colony PCR with *X. fastidiosa*-specific primers (RST31/RST33) and gene specific primers (pilA1-OFR- 405-F/pilA1-ORF-R). Complementation inserts were also confirmed by Sanger sequencing.

**Table 3.** Plasmids.

| Plasmid | Description | Source |
| --- | --- | --- |
| pCR8/GW/TOPO | Commercially available cloning vector with 3'-T overhangs. Compatible with Gateway® destination vectors, Sp<sup>R</sup> | Invitrogen |
| pCR8-ΔpilA1-chl | *pilA1* gene deletion construct containing chloramphenicol resistance marker flanked by ~1.5kb upstream and downstream sequences of *pilA1* gene, *pilA1* ORF is deleted, Sp<sup>R</sup>, Cm<sup>R</sup> | This study |
| pCR8-pilA1-ORF | *pilA1* complementation construct containing the *Xf pilA1* OFR and flanking regions. Used with Gateway pAX1-GW destination vector, Sp<sup>R</sup> | This study |
| pAX1-GW | Gateway® destination vector used for *Xf* chromosomal gene complementation into neutral location via homologous recombination, Cm<sup>R</sup>, Gm<sup>R</sup> | (45) |
| pAX1-pilA1-ORF | Gateway® complementation construct with ORF of *Xf pilA1* and flanking regions, Gm<sup>R</sup> | This study |

### Cell aggregation assay

*X. fastidiosa* strains were grown on PD3 agar plates and incubated at 28°C for 6-7 days. After incubation, bacteria cells were scraped off plates and resuspended in 5ml of liquid PD3 media (per sample) to OD600 = 0.10. Liquid cultures were grown in sterile 15 mL polypropylene test tubes at 28°C without shaking for 6-7 days. At least 3 replicates per strain were included. Cell aggregation was quantified using the OD600 of the upper culture (ODs) and the OD600 of the total culture (ODT). ODs, which is composed mostly of dispersed cells, was determined by measuring OD600 of undisturbed cultures. ODT was measured after aggregated cells were dispersed using a pipette and vortexing. The relative percentage of aggregated cells was estimated using the formula: [(ODt - ODs)/ODt] x100(46). The assay was repeated at least three separate times.

### Cell attachment assay

All procedures for setting up the attachment assays were performed aseptically. *X. fastidiosa* strains used in the attachment assays were grown on PD3 agar plates and incubated at 28°C for 6-7 days. After incubation, bacteria cells were scraped off plates and resuspended in 1 mL (per sample) of liquid PD3 medium. Small volumes of the concentrated cell suspensions were pipetted into 5ml (per sample) of fresh PD3 medium until a concentration of OD600 = 0.03-0.05 was reached. Aliquots of 100 µl of cell suspensions were added to individual wells of sterile 96-well polystyrene plates with lids (Nunclon, Cat #163320). Of the remaining cell suspension, 1 mL of each sample was transferred into sterile 1.5 mL centrifuge tubes and incubated at 28°C for 4 days (for gDNA extraction and qPCR later). Uninoculated liquid PD3 medium was used as a negative control. To minimize evaporation issues, we did not use wells from the outer most rows and columns of the plates. Plates were double wrapped with parafilm and incubated at 28°C for 4 days. Cell attachment was quantified using crystal violet staining. Media was removed from 96-well plates and the wells were washed three times with distilled water to remove unbound (planktonic) cells. Cells adhering to the sides of individual wells were stained with 100 μl of 0.1% (*w/v*) crystal violet for 25 min at room temperature. Crystal violet solution was removed from wells and wells were washed three times with distilled water. Crystal violet stain retained by attached cells was eluted by adding 100 μl of 30% acetic acid(47) to each well and quantified using a micro plate reader (Tecan Infinite M1000 PRO) at 550nm wavelength. OD550 results were normalized to CFU/ml of cells determined by qPCR. For qPCR, 1 ml aliquots of cells were centrifuged at 9000 rpm for three minutes to pellet cells, then frozen at -20°C until DNA extraction. DNA extraction was performed using a DNeasy Blood & Tissue Kit (Qiagen) following the manufacturer’s protocol for gram-negative bacteria. DNA was resuspended in 50 μl of sterile DEPC water (Invitrogen). One µl of each DNA sample was used as template for qPCR with Applied Biosystems PowerUp™ SYBR™ Green Master Mix (ThermoFisher) and primers targeting the *X. fastidiosa* chromosome (XfITS145-60F/ XfITS145-60R, Table 4). Concentration in CFU/ml was determined based on a standard curve of *X. fastidiosa* DNA extracted from samples with known CFU/ml concentrations. Cell attachment assays were repeated three separate times.

**Table 4.**
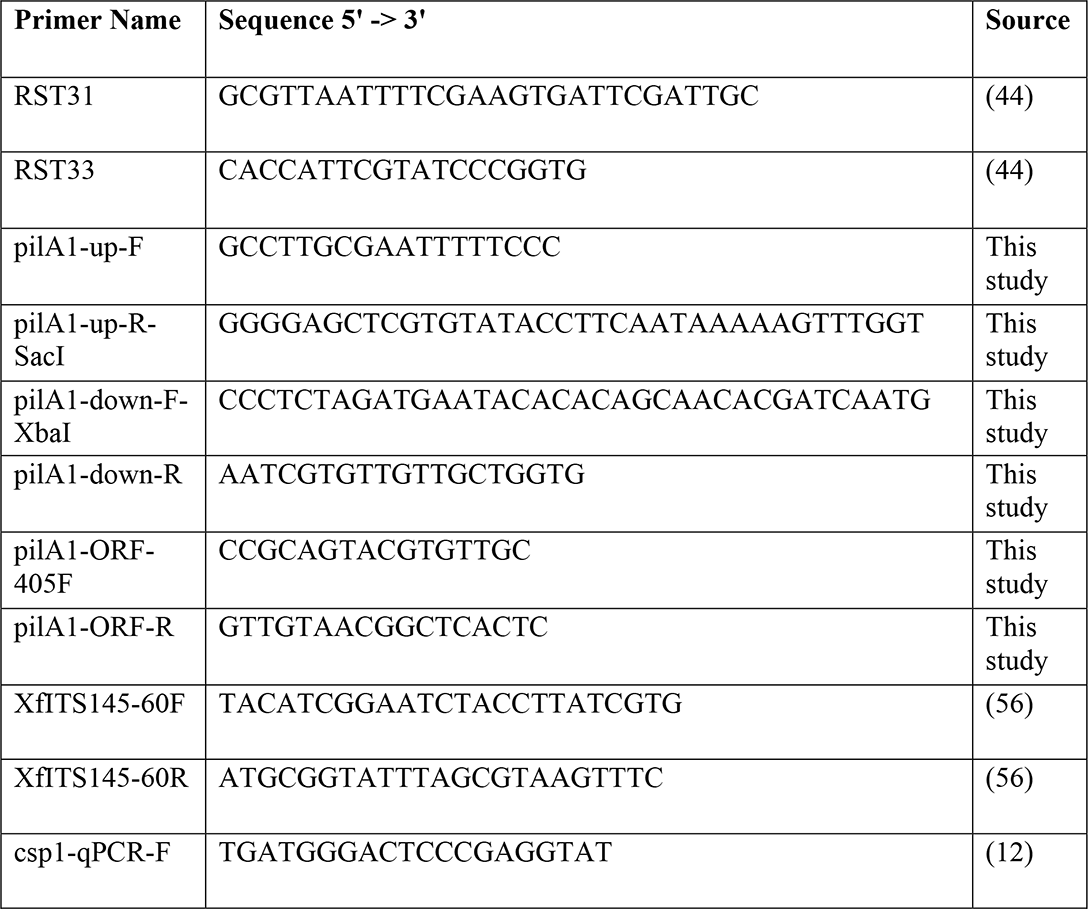
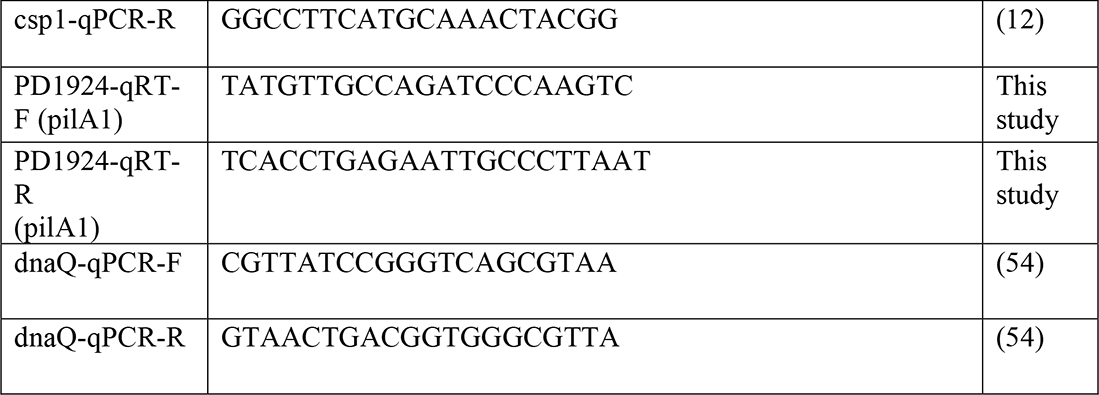
Primers

### Cell viability assay

Wild type, *Δcsp1*, and *Δcsp1/csp1+* strains were grown on PD3 agar plates for 7-13 days at 28°C. Cells were scraped off plates and resuspended in 1XPBS and diluted to OD600 = 0.01. One ml aliquots of each sample were reserved for gDNA extraction for cell quantification by qPCR. 90 µl of cell suspensions were added to individual wells of sterile 96-well plates and 10 µl of AlamarBlue Cell Viability reagent (Invitrogen) was mixed into each well. Plates were incubated in the dark at 37°C for 2 hours. Fluorescence was measured at 560 nm excitation/590 nm emission using a Tecan Infinite M1000 Pro plate reader. Cell aliquots reserved for qPCR were centrifuged at max speed for three minutes to pellet cells, then frozen at -20°C until DNA extraction. DNA extraction was performed using a DNeasy Blood & Tissue Kit (Qiagen) following the manufacturer’s protocol for gram-negative bacteria. DNA was resuspended in 50 µl of dH2O. One µl of each DNA sample was used as template for qPCR with Applied Biosystems PowerUp™ SYBR™ Green Master Mix (ThermoFisher) and primers targeting the *X. fastidiosa* chromosome (XfITS145-60F/ XfITS145-60R, Table 4). Concentration in CFU/ml was determined based on a standard curve of *X. fastidiosa* DNA extracted from samples with known CFU/ml concentrations. Fluorescence readings were normalized to CFU/ml.

### Transmission Electron Microscopy of *X. fastidiosa*

*X. fastidiosa* cells were grown on modified PW agar (omit phenol red and add 1.8 g/L of bovine serum albumin)(22) for two to three days. 3mm 300 mesh TEM grids were placed directly on bacteria cells growing on agar media for 5 seconds. Grids were immediately placed on a drop of 1.0% phosphotungstic acid for 30 seconds. Excess stain was wicked off the grid and placed in the FEI Helios Nanolab 650 SEM for STEM imaging.

### RNA-seq analysis

#### Bacteria growth conditions

WT Stag’s Leap and Δ*csp1* strains were grown on PD3 agar plates for 6 days at 28°C (6 plates per strain). Cells were aseptically harvested from each plate for all strains and immediately frozen on dry ice for RNA extraction later. Three replicates were included per strain, and each replicate sample included cells from two separate plates.

#### RNA preparation

Total bacterial RNA was extracted from frozen cells using the Trizol extraction method as described. Briefly, 1ml of Trizol (Invitrogen) reagent was added to each sample (in 1.5ml centrifuge tubes) and incubated at room temperature for 5 minutes. Samples were centrifuged to remove debris and the supernatant was transferred into new 1.5 ml tubes. 0.2 mL of chloroform was added to each sample and mixed thoroughly. Samples were centrifuged at 12,000 x g for 15 minutes. Following centrifugation, the colorless upper aqueous phase containing the RNA was transferred into a fresh tube and RNA was precipitated by adding 0.5 mL of room temperature isopropyl alcohol. Samples were incubated at room temperature for 10 minutes and centrifuged at 12,000 x g for 10 minutes. The RNA pellet was washed twice with 1 mL of 75% ethanol. Ethanol was removed and the RNA pellet was air dried and dissolved in DEPC-treated water. Total RNA was quantified using Quant-iT™ RNA Assay Kit (Invitrogen). 5ug of total RNA was treated with DNase I (Thermo Fisher) following the manufacturer’s protocol. DNase-treated RNA was re-precipitated using 0.1 volume sodium acetate and 3X volume ethanol. Poly(A) tail was added to the bacterial mRNA using Poly(A) Tailing Kit (Invitrogen) following the manufacturer’s protocol. RNA was re-precipitated using sodium acetate and resuspended in DEPC water (TE buffer was not used because EDTA concentrations as low as 1 mM will inhibit activity of exonuclease used in the next step). Ribosomal RNA was removed using the Lucigen Terminator 5’-Phosphate Dependent Exonuclease kit following the manufacturer’s protocol. The reaction was terminated, and the remaining RNA was precipitated using sodium acetate and ethanol. RNA quantity and quality were measured on the Agilent Bioanalyzer 2100 prior to cDNA library synthesis.

#### Nanopore cDNA library preparation

cDNA library synthesis was performed using the Oxford Nanopore direct cDNA synthesis kit (Oxford Nanopore) following the manufacturer’s protocol. All cDNA synthesis-specific reagents and consumables used were included in the kit unless stated otherwise. 250ng of PolyA+ mRNA resuspended in 7.5 μl of nuclease-free water (per sample) was added to DNA LoBind tubes and centrifuged briefly. Reverse transcription and strand-switching were performed by first adding 2.5 μl VNP primer and 1 μl 10mM dNTPs (ThermoFisher) to each mRNA sample and incubating at 65°C for 5 minutes, followed by immediately cooling on ice. In separate tubes, 4 μl of 5X Maxima H Minus RT Buffer (Thermo Fisher), 1 μl RNaseOUT (ThermoFisher), 1 μl Nuclease-free water, and 2 μl Strand-Switching Primer (SSP) were combined and added to the mRNA samples. Samples were incubated at 42°C for 2 minutes, after which 1 µl of Maxima H Minus Reverse Transcriptase (Thermo Fisher) was added each sample. Samples were incubated at 42°C for 90 minutes, followed by heat inactivation of reaction at 85°C for 5 minutes. Residual RNA was digested by adding 1 µl of RNase Cocktail Enzyme Mix (ThermoFisher) to each reverse transcription reaction. Samples were transferred to new 1.5 mL DNA LoBind Eppendorf tubes and cDNA purified using 17 µl of resuspended AMPure XP beads (Agencourt) flowing the manufacturer’s protocols. Samples mixed with AMPure XP beads were centrifuged briefly and beads (bound to cDNA) were immobilized to tube walls using a magnetic tube rack. Tubes were kept on the magnetic rack and the supernatant removed and discarded. The beads were washed twice with 200 µl freshly prepared 70% molecular grade ethanol. Residual ethanol was removed, and bead pellets were air dried briefly (not to the point of the pellet cracking). Tubes were removed from the magnetic rack and bead pellets were resuspend in 20 µl of DEPC water (Invitrogen). Tubes were put back on the magnetic rack to separate the eluate from the AMPure XP beads, and 20 µl of eluate from each sample were transferred into separate 0.2 mL PCR tubes for cDNA second strand synthesis. 25 μl 2X LongAmp Taq Master Mix (New England Biolabs), 2 μl PR2 primer, and 3 μl of DEPC water was added to each 20 µl eluate sample. Thermal cycler conditions were as follows: 94°C for 1 min, 50°C for 1 min, 65°C for 15 mins, hold at 4°C until next step. Samples were transferred into 1.5 mL DNA LoBind tubes and cDNA purified using 40 µl of AMPure XP beads following the same protocol as previously described. Purified cDNA was eluted in 21 µl of DEPC water. 1 µl of purified cDNA was analyzed on the Agilent Bioanalyzer 2100 to check the quality and quantity. End repair and dA-tailing of fragmented cDNA were performed by mixing: 20 µl cDNA sample, 30 µl Nuclease-free water, 7µl Ultra II End-prep reaction buffer (New England Biolabs), and 3 µl Ultra II End-prep enzyme mix (New England Biolabs). Samples were incubated at 20°C for 5 minutes, followed by 65°C for 5 minutes in a thermal cycler. Samples were transferred into 1.5 mL DNA LoBind Eppendorf tubes and purified using 60 μl of AMPure XP beads following the same protocol as before. Samples were resuspended in 22.5 µl of DEPC water and transferred into a clean 1.5 ml Eppendorf DNA LoBind tubes.

#### Barcode Ligation

Individual cDNA libraries (12 total) were ligated with unique native barcodes (Oxford Nanopore Native Barcode Expansion set 1-12) following the manufacturer’s protocols. 22.5 µl of cDNA was combined with 2.5 µl native barcode and 25 µl Blunt/TA Ligase Master Mix (New England Biolabs). Samples were incubated at room temperature for 10 minutes, and barcoded cDNA libraries were purified using 40 μl of AMPure XP beads and resuspended in 26 µl DEPC water following the same protocol as used during cDNA library preparation. 1 µl of each sample was quantified using the Quant-iT High-Sensitivity dsDNA Assay Kit (Thermo Fisher). The quantity of cDNA for one replicate sample of WT 28°C was too low and was excluded from further experiments. The barcoded cDNA libraries from the remaining 11 samples were pooled in equal ratios to obtain 700 ng total DNA and final volume adjusted to 50 μl using nuclease free water and loaded into the Nanopore MinION flow cell (FLO-MIN106). The sequencing reaction was run for 23 hours and generated approximately 4.04 million total reads in FAST5 format.

#### Data Analysis

Nanopore FAST5 files were converted into FASTQ files using the basecalling program Guppy(48) . Barcoded samples were demultiplexed using Deepbinner(49). The program Porechop was used to trim off adapter sequences from the demultiplexed FASTQ reads. The *X. fastidiosa* Temecula-1 cDNA reference transcriptome (ftp://ftp.ensemblgenomes.org/pub/bacteria/release-44/fasta/bacteria_18_collection/xylella_fastidiosa_temecula1/cdna/) was indexed and FASTQ reads were mapped to the reference using Minimap2(50). After mapping, aligned reads were quantified using Salmon(51). A table summarizing transcript-level estimates for use in differential gene analysis was created using the R package Tximport(52). Differential expression analysis was performed using the R package DESeq2(53). Descriptions of the programs used and web addresses for downloading the source codes are listed in Supplemental Table 2 (S2).

### qRT-PCR Gene Expression Analysis

Quantitative reverse transcriptase PCR (qRT-PCR) was used to confirm gene expression results of several differentially expressed genes of interest from the RNA-Seq experiment, as well as monitor expression of *csp1* during different *X. fastidiosa* growth stages. For RNA extraction to confirm differentially expressed genes, cells were grown under the same conditions as for the RNA-Seq experiment. For *csp1* expression, cells were grown as described in the cell viability assay. Total RNA was extracted as described in the RNA-Seq methods section using the Trizol (Invitrogen) method. gDNA was removed using Baseline Zero DNase (Lucigen) following the manufacturer’s protocols and RNA reprecipitated using 0.1 volume of sodium acetate and 2-3 volumes 100% ethanol. Purified RNA was quantified using a Quant-IT RNA Assay kit (Thermo Fisher Scientific). Removal of residual DNA was confirmed by DNA-specific quantification using a Quant-IT dsDNA Broad Range Assay Kit (Thermo Fisher Scientific). For cDNA synthesis, 500 ng of total RNA was reverse transcribed with random primers using an iScript gClear cDNA synthesis kit (BioRad) and including a no-RT and controls for each sample. 1 µl of each cDNA sample was used as template for qPCR using Applied Biosystems PowerUp SYBR Green Master Mix (Thermo Fisher Scientific). *X. fastidiosa dnaQ* gene, which is a stable reference gene in *X. fastidiosa*(54), was used to normalize expression of other target genes. Primer sequences for target genes are listed in Table 4. PCR cycling conditions were based on recommended protocol provided by PowerUp SYBR Green Master Mix and the melting temperature of the different primer sets. Experiments were repeated three independent times and relative gene expression was calculated with BioRad CFX Manager software.

### Plant Virulence Assays

#### Plant inoculations

Wild type Stag’s Leap, Δ*pilA1*, and Δ*pilA1/pilA1*+ strains were grown on PD3 agar plates for 5-7 days and then scraped off plates and resuspended in 1XPBS at concentration OD600=0.25 (∼1x10^8^ CFU/mL). Susceptible (cv Chardonnay) one-year-old potted grapevines were inoculated using a pinprick inoculation method(55). Mock inoculations using 1XPBS were used as negative controls. 20 plants were inoculated with wild type, 15 plants with Δ*pilA1*, 15 plants with Δ*pilA1/pilA1*+, and 10 plants with 1XPBS. Plants were labeled with number codes and placed randomly within a climate-controlled greenhouse. The plants were monitored weekly for development of scorching symptoms. Once disease symptoms began to develop (5 weeks post inoculation for this experiment), plants were given a disease index score between 0-5 based on a rating scale previously developed(55). A disease score of 0 indicates no disease symptoms and a score of 5 represents severe disease symptoms and plant death. Representative images of disease ratings were provided courtesy of Yaneth Barreto-Zavala and included in Supplemental Figure 2 (S2). Plants were rated until 12 weeks post-inoculation. Area under the disease progress curve (AUDPC) was calculated using average disease intensity over time (weeks) with the Agricolae package for R (https://CRAN.R-project.org/package=agricolae). Plant infection assays were conducted during June-September 2020

#### qPCR Quantification of Bacterial Populations

At 9- and 12-weeks post-inoculation, petiole samples from infected and mock-inoculated plants were collected for DNA extraction and qPCR quantification of *in planta* bacterial populations. 2-3 petiole samples from each plant were pooled, and samples were lyophilized using a FreeZone (LABCONCO) freezer dryer at -80°C for 24-48 hours. Lyophilized samples were pulverized with 3mm Tungsten Carbide beads (Qiagen) using a Tissue Lyser II (Qiagen) for a total of 4 minutes at 30 r/s. One mL of DNA extraction buffer (20mM EDTA, 350mM Sorbitol, 100mM Tris HCL) with 2.5 % polyvinylpyrrolidone was added to each sample and centrifuged at 14,000 rpm for 5 minutes. All centrifugation steps were performed at 4°C. Supernatant was removed, and pellet was washed with an additional 1 mL of DNA extraction buffer and centrifuged for 10 minutes at 14,000 rpm. Supernatant was removed and pellet was re-suspended with 300 μL of DNA extraction buffer, 300 μL of lysis buffer (50Mm EDTA, 2M NaCl, 2% CTAB, 200mM Tris HCl) and 200 µL of 5% sarcosyl. Tubes were incubated for 45 minutes at 65°C and mixed by vortexing every 15 minutes. After incubation 700 μL of chloroform:isoamly alcohol (24:1) was added to each tube and inverted to mix samples. Samples were then centrifuged at 9500 rpm for 5 minutes. The upper phase was transferred to a new tube and 800 μL of phenol:chloroform:isoamly alcohol (25:24:1) was added. Samples were mixed and centrifuged at 9500 rpm for 5 minutes. The upper phase was transferred to a new tube and 1 mL of isopropanol was added to precipitate the DNA. Samples were mixed and centrifuged at 12000 rpm for 15 minutes. Supernatant was removed and pellet was washed with 300 µL of chilled 70% ethanol and dried under a fume hood. DNA was then re-suspended in 50 μL of TE buffer. Samples were diluted 1:10 in sterile dH2O prior to quantification by qPCR.

For qPCR, 5 µl of DNA was used as template with Applied Biosystems Fast SYBR Green Master Mix and primers targeting the *X. fastidiosa* chromosome (XfITS145-60F/ XfITS145-60R, Table 4). A standard curve for quantification was made with 10-fold dilutions of *X. fastidiosa* DNA extracted from 1x10^8^ CFL/mL cell suspension combined with uninfected grape DNA in a 2:1 ratio. PCR consisted of 95°C for 3 minutes followed by 35 cycles of 95°C for 30 seconds and 60°C for 30 seconds. PCR was performed using a BioRad CFX96 instrument. CFU/ml as determined by qPCR was normalized to total DNA concentration in ng/µl. Total DNA concentration of original samples was determined using Quant-iT™ dsDNA Assay Kit (Thermo Fisher).

## Acknowledgments

We would like to thank Brandon Ortega and Nathaniel Luna for technical support and Yaneth Barreto-Zavala for providing the images used for the grapevine symptoms rating scale. We would also like to thank Leonardo De La Fuente and Marcus Merfa for helpful suggestions regarding imaging pili. Funding for this work was from United States Department of Agriculture (USDA) Agricultural Research Service appropriated project 2034-22000-012-00D. Mention of trade names or commercial products in this publication is solely for the purpose of providing specific information and does not constitute endorsement by USDA. USDA is an equal opportunity provider and employer.

